# Semi-Automated Identification of EKG and Trigger Artifacts in EEG Using ICA and Spectral Characteristics

**DOI:** 10.64898/2026.04.08.717297

**Authors:** Amilcar J. Malave, Blair Kaneshiro

## Abstract

A persistent bottleneck in post-Independent Component Analysis (ICA) Electroencephalogram (EEG) preprocessing is the manual identification of artifact components for removal. In practice, this step can be slow, subjective, and difficult to standardize, particularly for cardiac contamination and trigger-related leakage, where artifact structure may be distributed across multiple components or appear outside the highest-variance Independent Components (ICs). We developed the *SENSI-EEG-Preproc-ICA-EKG-Trigger Module* to make this stage faster and more reproducible without removing the user from the decision process. The Module is a semi-automated MATLAB framework for post-ICA screening of cardiac and trigger-related artifact components using spectral characteristics. EKG candidates are prioritized by detecting harmonic structure around a physiologically plausible heart-rate fundamental, whereas trigger-related candidates are prioritized by measuring harmonic concentration at frequencies determined by the known repetition period of the trigger sequence. The resulting candidates are then reviewed in dedicated interfaces that present scalp topography, time-domain activity, and frequency-domain structure together, allowing the final classification to be confirmed or corrected by the user. In this way, the Module narrows the search space while preserving interpretability and explicit human control over the final keep/remove decision. The release includes a public codebase, a user manual, example workflows, and an accompanying example dataset. This paper presents the Module as a practical methods-and-software contribution for post-ICA EEG cleaning.

## Introduction

Electroencephalography (EEG) recordings are routinely contaminated by non-neural activity arising from ocular motion, muscle activity, electrode instability, environmental interference, and physiological sources such as cardiac activity. Independent component analysis (ICA) has therefore become a standard element of many EEG preprocessing pipelines because it can decompose multichannel recordings into statistically independent components that often isolate distinct neural and artifactual processes, allowing selective component rejection before reconstruction of the cleaned sensor-level data [1–3]. In practice, however, obtaining an ICA decomposition does not eliminate the main interpretive challenge: the user must still decide which independent components reflect artifact and which should be preserved.

In many ICA-based workflows, this decision is guided by the joint inspection of component scalp projections, activation time courses, and spectral profiles [3]. A number of tools have been developed to support this stage, including semi-automatic review aids and trained independent-component classifiers such as CORRMAP, SASICA, and ICLabel [3–5]. Here, we introduce an artifact-specific Module for two contamination types that benefit from targeted scoring rules and a streamlined review workflow. Depending on their needs, users can decide which main function(s) they wish to call (i.e., which contamination type(s) they wish to address).

One artifact family considered here is cardiac contamination, or electrocardiogram (EKG). Current approaches for cardiac artifact handling already include ICA-based removal, often aided by a recorded EKG channel or QRS timing, as well as template-based subtraction, regression or adaptive filtering, and projection or other subspace-based approaches [6– 9]. The narrower contribution of the present Module is a practical *post-ICA* screening procedure that rapidly scores all selected components for EKG-like spectral structure associated with quasi-periodic cardiac activity, ranks candidates for inspection, and then leaves the final rejection decision to the user through a dedicated review interface. This emphasis on screening all components is intentional, since relevant artifact may remain visible in higher-numbered (i.e., lower-variance) components even when it is not confined to the most visually dominant part of the decomposition.

The second artifact family addressed here is contamination associated with digital event triggers, referred to throughout this preprint as *Digital Input (DIN)* artifact. In the present workflow, DIN events are used to mark stimulus timing, and in Electrical Geodesics, Inc. (EGI)/Net Station-style acquisitions they may be conveyed to the amplifier as TTL-like 5V digital pulses connected to the system’s digital inputs [10]. In some recording setups, these trigger pulses, or their electrical coupling through cabling and grounding, may leak into the EEG acquisition. In the affected EEG channels, this contamination may appear as sharp event-locked transients in the time domain and, when the triggering repeats at a fixed period, as comb-like harmonic structure in the frequency domain. The DIN detector introduced here targets this specific leakage scenario by using the known trigger period to quantify whether the an ICA component’s spectrum contains energy concentrated at the expected harmonic frequencies.

We present a practical semi-automated MATLAB Module for rapid identification of EKG- and DIN-related independent components after ICA decomposition. For EKG detection, the method searches each component for a physiologically plausible low-frequency fundamental and evaluates whether the corresponding spectrum exhibits harmonic structure consistent with cardiac contamination. For DIN detection, the method uses the known trigger repetition period to compute a harmonic-energy score that contrasts power near expected harmonics against the surrounding baseline spectrum. In both cases, the score is used to prioritize components for review, not to replace final user judgment.

An equally important design choice is the retention of an explicit review stage. Final classification is performed through dedicated interfaces that display, for each candidate component, its scalp map, a representative time-domain segment, and its spectral profile. The intent is to accelerate component review while preserving transparency and user control over the final rejection decision. The Module should therefore be viewed as a decision-support tool tailored to two artifact classes, not as a universal black-box artifact classifier.

The intended operating point is after filtering, bad-channel handling, and ICA decomposition have already been completed. The DIN detector further assumes that the relevant trigger period is known, and the EKG detector assumes that cardiac contamination is sufficiently represented in the component activations to be detectable by the proposed score. This preprint is thus a methods-and-software contribution with representative examples. The broader codebase release includes three main components:

1. A publicly available GitHub repository containing the complete codebase, documentation, and usage examples [11].^1^
2. A standalone example dataset used for the illustrative analyses, with accompanying scripts for automated download and local setup [12].^2^
3. This preprint, which describes the Module’s contents, algorithms, and user experience.

This Module has been developed as part of a larger EEG preprocessing pipeline (in preparation) that handles identification of electrooculogram (EOG) artifacts through a separate post-ICA decision process, namely on the basis of component activation correlations with precomputed EOG channels as well as visual inspection (as in e.g., [13]). During development of the present Module, we also explored topography-based and direct time-course criteria. These were not retained as the primary automated rules because they proved less robust than the final frequency-based scoring approach for the recordings examined here, but descriptions of these alternative strategies and the location of example code in the repository are provided in the User Manual.

The remainder of this preprint is organized as follows. We first describe the Module overview and its intended use within an EEG preprocessing workflow. We then summarize the codebase structure and present the EKG and DIN identification procedures, the associated review interfaces, and the alternative criteria considered during development. Representative examples are then used to illustrate the workflow in practice. We conclude with a discussion of strengths, limitations, and directions for future validation and extension.

### Codebase

#### Code access and license

The SENSI-EEG-Preproc-ICAEKG-Trigger Module is distributed through a public GitHub repository.^1^ The present preprint documents the current tagged release of the Module. The software is released under the MIT license.

Alongside the codebase, the release includes a standalone example dataset hosted on the Stanford Digital Repository (SDR).^2^ The example script in the repository, `example.m`, automatically retrieves the example data files and places them in the local ***ExampleData*** folder.

If using the Module (code, figures, or methods), please cite (i) the GitHub repository, (ii) this preprint, and (iii) the example dataset when used for demonstrations or external projects

#### Dependencies

The Module requires MATLAB and depends on two MathWorks toolboxes: Signal Processing Toolbox and Statistics and Machine Learning Toolbox. Development and testing were performed in MATLAB R2024b. The current release was validated on both Windows and Linux systems.

#### Folders and files in the repository

#### ExampleData

By default, contains only a `readme.txt` file; due to file size, example datasets used by `example.m` are not included directly in the GitHub repository. Instead, `example.m` downloads the required files from the SDR into this folder.

#### Figures

Default output folder used by `example.m` for saving generated figures.

#### HelperFunctions

Contains helper functions called internally by the automated detectors and related plotting routines. These functions are not typically called directly by the user.

#### Sensor Layouts

Contains electrode-location files used for scalp topography visualization.

#### example.m

A complete walkthrough script showing how to configure the Module, download example data, transform the recording into ICA space, run `autoDetectDIN()`and `autoDetectEkg()`, and inspect outputs through the interactive review interfaces. This is the recommended starting point for new users.

#### autoDetectDIN.m

Main user-called function for identifying DIN-contaminated ICA components. It computes a harmonic-based score using the known trigger repetition period and returns candidate components for review.

#### autoDetectEkg.m

Main user-called function for identifying EKG-contaminated ICA components. It searches for a physiologically plausible cardiac fundamental and associated harmonic structure, and returns candidate components for review.

#### reviewDINArtifactUI.m

Interactive review interface for DIN candidates. It displays suspected components together with scalp maps, time-domain activity, and frequency-domain views to support final user confirmation.

#### reviewEkgArtifactUI.m

Interactive review interface for EKG candidates. It similarly presents scalp, time-domain, and spectral views for final confirmation of candidate artifact components.

#### README.md

A brief overview of the Module, including expected inputs/outputs and pointers to documentation.

#### LICENSE

License for the repository (MIT).

#### User Manual.pdf

The User Manual for the current release, including background, setup instructions, a full walkthrough of `example.m`, interpretation guidance, and detailed documentation of main and helper functions.

#### .gitignore

Git configuration file (no user action required).

The current release is therefore distributed as a self-contained and documented MATLAB Module, with separate components for software, example data, and written documentation. This structure is intended to support transparent reuse, reproducible demonstration of the workflow, and clear separation between the automated scoring routines and the final user-guided review stage.

### Functionalities of the Module

The SENSI-EEG-Preproc-ICA-EKG-Trigger Module is designed around two artifact-specific post-ICA detection procedures: one for cardiac (EKG) contamination and one for DIN-related trigger leakage. In both cases, the workflow follows the same general logic: ICA components are first screened automatically using a frequency-domain metric tailored to the artifact family, and the resulting candidates are then passed to an interactive review interface for final confirmation.

#### EKG artifact detection

The main entry point for EKG detection is `autoDetectEkg()`, which operates on ICA component time series:

~~~
    [ekgSrc, ekgSus, metrics]=
       autoDetectEkg(*xICA, f s, nSources, opts*)
~~~

The function scans the selected ICA components and scores each one using a harmonic local signal-to-noise ratio (SNR) metric. The rationale is that cardiac contamination often produces a physiologically plausible low-frequency fundamental together with harmonics extending across the spectrum.

The purpose of the automatic stage is therefore to identify components whose spectra are unusually consistent with this pattern.

For each ICA component *c*, the time series *x*_*c*_(*t*) is converted to the frequency domain using Welch’s method:

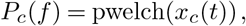

where *P*_*c*_(*f*) denotes the one-sided power spectral density (PSD). The Welch window length and overlap are controlled by the options structure `opts` (structure with detector parameters). If enabled, the log–log spectrum is linearly detrended over a user-defined frequency range before scoring in order to reduce broadband 1*/f* -like structure and emphasize narrowband periodic peaks.

The first detection step is to identify a candidate cardiac fundamental within a physiological search band:

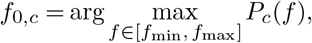

where the default range is *f*_min_ = 0.8 Hz to *f*_max_ = 2.0 Hz. This band is intended to capture plausible heart-rate fundamentals while excluding unrelated low-frequency structure. Given the detected fundamental *f*_0,*c*_, expected harmonic frequencies are defined as

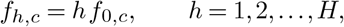

where *H* is the maximum number of harmonics considered. For each harmonic *f*_*h,c*_, two local frequency regions are defined. The first is a narrow peak window centered at the harmonic,

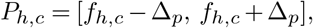

where Δ_*p*_ is a small half-width in Hz. The second is a surrounding neighborhood used to estimate the local spectral baseline,

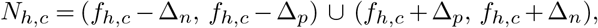

with Δ_*n*_ *>* Δ_*p*_.

The local harmonic SNR is then defined as the ratio between harmonic peak power and its surrounding baseline:

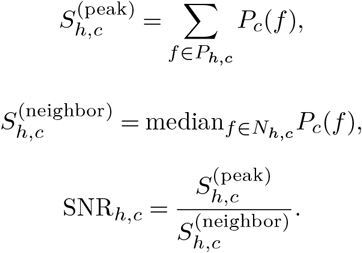

Large values indicate that the harmonic stands out clearly relative to nearby frequencies. This local contrast makes the detector less dependent on absolute power and more sensitive to structured harmonic peaks.

To summarize harmonic structure across frequencies, the harmonic SNR values are aggregated into a composite score. Let *v*_*c*_ be the harmonic composite score for component *c*:

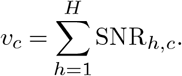

The intuition is that true cardiac contamination should not only contain a single peak in the heart-rate band, but a broader harmonic series. Summing across harmonics therefore increases robustness to variability in any single harmonic.

The resulting composite scores are then standardized across the scanned components using a robust median/MAD transformation.

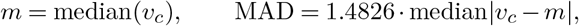

and the robust score is

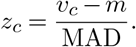

These values are used to produce two ranked output lists. Components with large outlier values (*z*_*c*_ *>* 5 in the current implementation) are returned as strong EKG candidates in `ekgSrc`. If fewer than three such components are found, the remaining slots are filled by the next highest-ranked components with more moderate scores (*z*_*c*_ *>* 3), which are returned as `ekgSus`. The function therefore returns at most three total components across both lists.

The final EKG decision is made through `reviewEkgArtifactUI()`, which presents to the user each candidate component using the same three complementary views commonly used in ICA interpretation: scalp topography, a representative time-domain segment, and the full-series magnitude spectrum. Components initially flagged as EKG candidates are shown in red, and the user can toggle each component between *EKG/remove* and *keep*.

### DIN artifact detection

The main entry point for DIN detection is `autoDetectDIN()`, which operates on ICA component time series in the same post-ICA setting:

~~~
    [dinSrc, info] =
       autoDetectDIN(*xICA, f s, nSources, dinOpts*)
~~~

The DIN detector uses the known trigger repetition period to evaluate whether a component exhibits spectral energy concentrated at the expected harmonics of the digital trigger sequence.

Let *x*_*c*_(*t*) denote the ICA time series for component *c*. The first step is spectral estimation using Welch’s method:

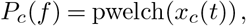

where *P*_*c*_(*f*) is the one-sided PSD. The DIN metric is parameterized by the DIN repetition period *T*_DIN_ (in seconds), provided through the options structure. The corresponding base frequency is

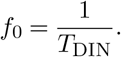

Expected harmonic frequencies are then defined as

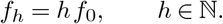

Only harmonics that fall within the valid analysis band are retained:

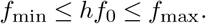

This yields the harmonic index set

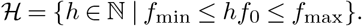

For each harmonic *f*_*h*_, a narrow peak window is defined:

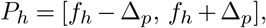

where Δ_*p*_ is a small half-width in Hz. The total harmonic energy is computed as

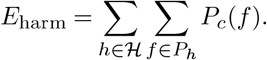

To estimate the local baseline around these harmonics, a local spectral neighborhood is defined for each harmonic:

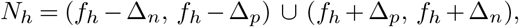

with Δ_*n*_ *>* Δ_*p*_. The baseline energy is then computed as

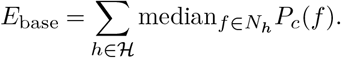

Using the median provides a robust estimate of local background power while limiting the influence of isolated peaks in the surrounding spectrum.

The DIN score for component *c* is defined as the harmonic energy ratio

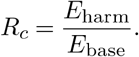

Large values of *R*_*c*_ indicate that spectral energy is concentrated disproportionately at the expected trigger-locked harmonics, consistent with structured digital interference.

These harmonic-ratio values are then interpreted relatively across the scanned ICA components. Let *S* denote the set of scanned components. The DIN metric is standardized as

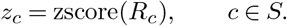

Positive outliers in this space are treated as likely DIN candidates. Candidate selection is carried out as a two-stage process with a hard cap of three total components. In the first stage, a fixed threshold *τ* = 2.5 is applied:

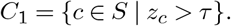

Components in *C*_1_ are ranked in descending order of *z*_*c*_, and up to three are assigned directly to `dinSrc`. In the second stage, if fewer than three components exceed the primary threshold, additional high-ranking components are added after re-standardization so that they can still be examined during review.

The final DIN decision is made through `reviewDINArtifactUI()`, which presents to the user each candidate component using the same triad of views used in EKG review: scalp topography, a representative time-domain segment, and the full-series magnitude spectrum. Components initially flagged as DIN candidates are shown in red, and the user can toggle each one between *DIN/remove* and *keep*.

### Representative ICA signatures of EKG and DIN artifacts

Before presenting the illustrative workflow, it is useful to show how these artifact classes typically appear in ICA space. Throughout the Module, candidate components are evaluated using three complementary views: scalp topography, a representative time-domain segment, and the frequency-domain representation.

### Representative EKG ICA component

A representative EKG-related ICA component is shown in Figure 1. In the time domain, cardiac contamination typically appears as a repeating beat-like waveform, often with sharp transient structure occurring at a roughly regular cadence. In the frequency domain, the defining feature is a physiologically plausible low-frequency fundamental, typically within the heart-rate range, together with clear integer harmonics. The scalp topography often appears as a broadly distributed pattern with a smooth polarity gradient (i.e., a dipole-like map). This is consistent with a far-field generator whose electric field propagates through a large volume conductor before being measured at EEG electrodes [14]. In practice, however, the exact orientation and distribution can vary across recordings, and cardiac activity is not always isolated to a single ICA component.

**Fig. 1.**
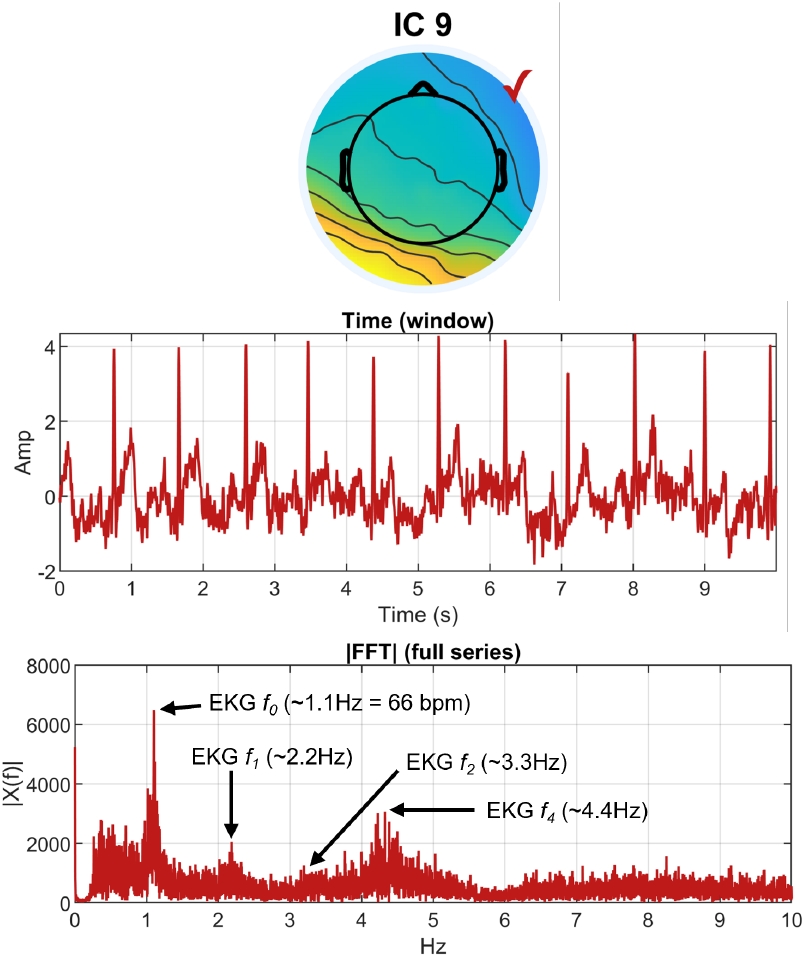
Representative EKG ICA component. Example of an ICA component showing the typical EKG pattern across the three review panels: a broad scalp projection, a repeating cardiac waveform in the time domain, and a low-frequency fundamental with harmonics in the spectrum. For display purposes, only 10 seconds of time-domain data are shown here, but the component topography and frequency-domain plot reflect the full data record (approximately 7 minutes).

### Representative DIN ICA component

A representative DIN-related ICA component is shown in Figure 2. In the time domain, it typically appears as brief sharp transients occurring at fixed intervals (in this case, 1 Hz). In the frequency domain, the main cue is a pronounced harmonic comb, with energy concentrated at the trigger repetition frequency and its harmonics. The scalp projection is often focal or hotspotlike, frequently near posterior midline electrodes such as Pz.

**Fig. 2.**
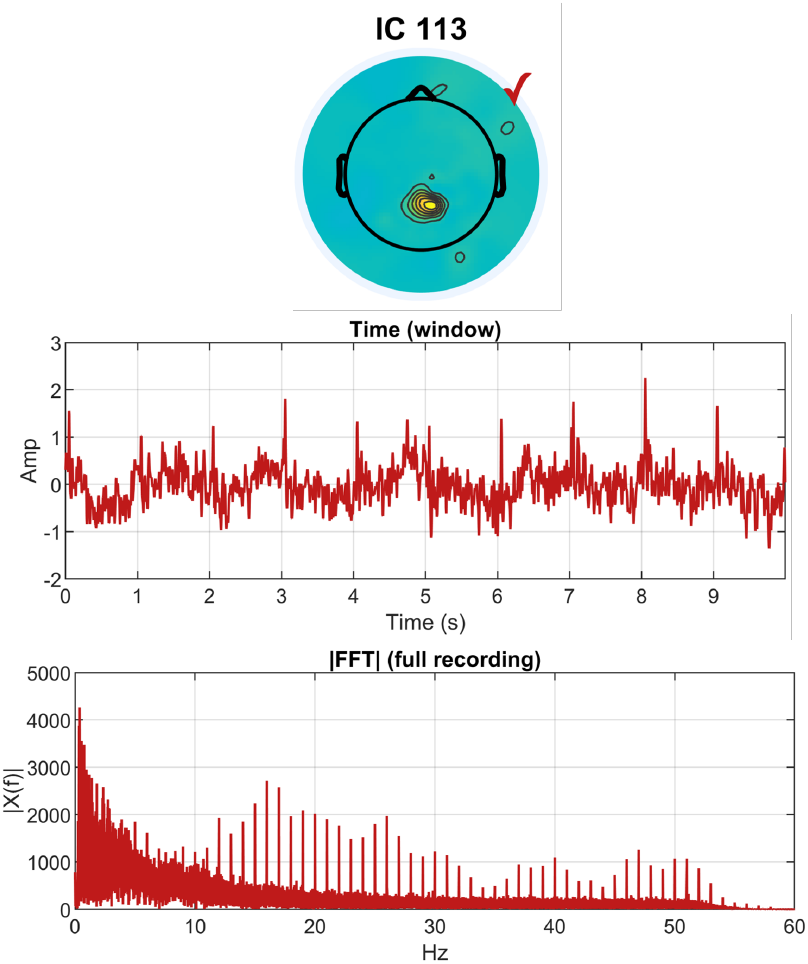
Representative DIN ICA component. Example of an ICA component contaminated by trigger-related leakage, showing a focal scalp projection, sharp recurring transients in the time domain, and a comb-like harmonic pattern in the spectrum. For display purposes, only 10 seconds of time-domain data are shown here, but the component topography and frequency-domain plot reflect the full data record (approximately 7 minutes).

### Illustrative analysis

This illustrative analysis demonstrates the user-facing workflow of the SENSI-EEG-Preproc-ICA-EKG-Trigger Module: obtaining example data, transforming the recording into ICA space, running the two automated detection procedures, and interpreting the interactive review outputs that lead to the final artifact-component list. The goal of this section is to show what users will encounter when running the Module and how the outputs should be interpreted at a high level. Detailed parameter descriptions, UI controls, and function-level documentation are provided in the User Manual.

#### Running the Module

The GitHub repository includes a runnable script, `example.m`, which demonstrates the complete analysis sequence.

#### Example data and preprocessing state

The example data are distributed separately through the Stanford Digital Repository (SDR) and are downloaded automatically into the local ***ExampleData*** folder by `example.m`. Each example file contains, at minimum, a filtered sensor-space data matrix **x**_Raw_, a sampling rate *f*_*s*_, and a list of bad channels whose rows have already been removed from **x**_Raw_. The illustrative workflow therefore begins from data that have already undergone basic filtering and bad-channel handling, consistent with the intended operating point of the Module.

The script next loads the electrode-location file used for scalp topography visualization and removes the entries corresponding to channels listed in `badCh`. It then obtains the ICA unmixing matrix **W**, either by loading a cached version from disk or by computing it directly from the current data if no cached file is available. The sensor-space recording is then mapped into ICA space according to

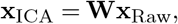

yielding the ICA components used by the automated detectors.

#### Automated detection and interactive review

After the data have been transformed into ICA space, the illustrative workflow proceeds in two steps: automated candidate selection followed by interactive confirmation. The example script first runs the DIN detector and launches the DIN review interface, and then runs the EKG detector and launches the corresponding EKG review interface. In both cases, the automated procedure reduces the search space to a small set of candidate components, while the final keep/remove decision is made by the user. Further details on configurable parameters and function-specific behavior are provided in the User Manual.

#### DIN review

The DIN portion of the workflow begins with `autoDetectDIN()`, followed by `reviewDINArtifactUI()`. The DIN review interface is shown in Figure 3. Each row corresponds to one ICA component and displays three complementary views: scalp topography, a representative time-domain segment, and the frequency-domain representation. In this example, Component 113 is initially flagged as a DIN artifact, while Components 48 and 5 are presented as additional review candidates. Based on the combined evidence across the three panels, only Component 113 shows the characteristic DIN pattern, so the default labeling is retained: Component 113 is flagged for removal, while Components 48 and 5 are kept.

**Fig. 3.**
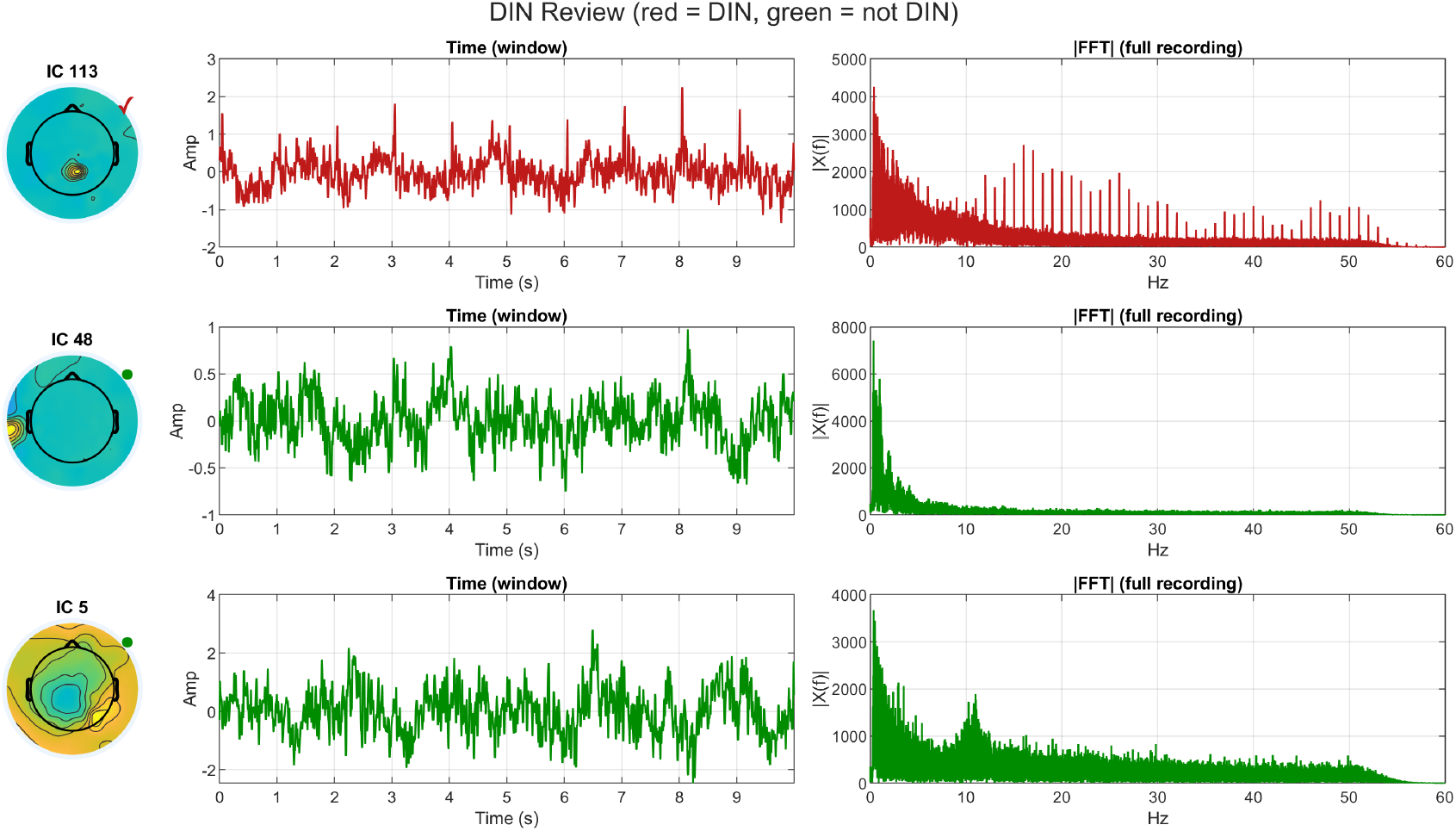
DIN interactive review interface. Component 113 is flagged for removal, while Components 48 and 5 are shown as additional review candidates. The final decision retains the default classification: Component 113 is removed and Components 48 and 5 are kept.

#### EKG review

The EKG portion of the workflow then begins with `autoDetectEkg()`,followed by `reviewEkgArtifactUI()`. The EKG review interfaces before and after user confirmation are shown in Figure 4 and Figure 5, respectively. As in the DIN review, each row presents the scalp map, a time-domain segment, and a frequency-domain view for one ICA component. In this example, Components 9, 33, and 48 are initially marked as EKG candidates. The time-domain plot for Component 9 shows the clearest cardiac pattern across all three views. Component 33 is less obvious in the displayed time segment, but its spectral structure remains consistent with cardiac contamination and supports removal. This is not unexpected, since cardiac activity is not always captured by a single ICA component. Different portions of the cardiac cycle, such as sharp QRS-like deflections versus slower wave components, may separate into different ICs. Component 33 may therefore reflect a slower cardiac-related contribution rather than the sharper transient structure seen more clearly in Component 9. Component 48 shows weaker evidence than the other two. After review, Components 9 and 33 remain marked for removal, whereas Component 48 is toggled to keep.

**Fig. 4.**
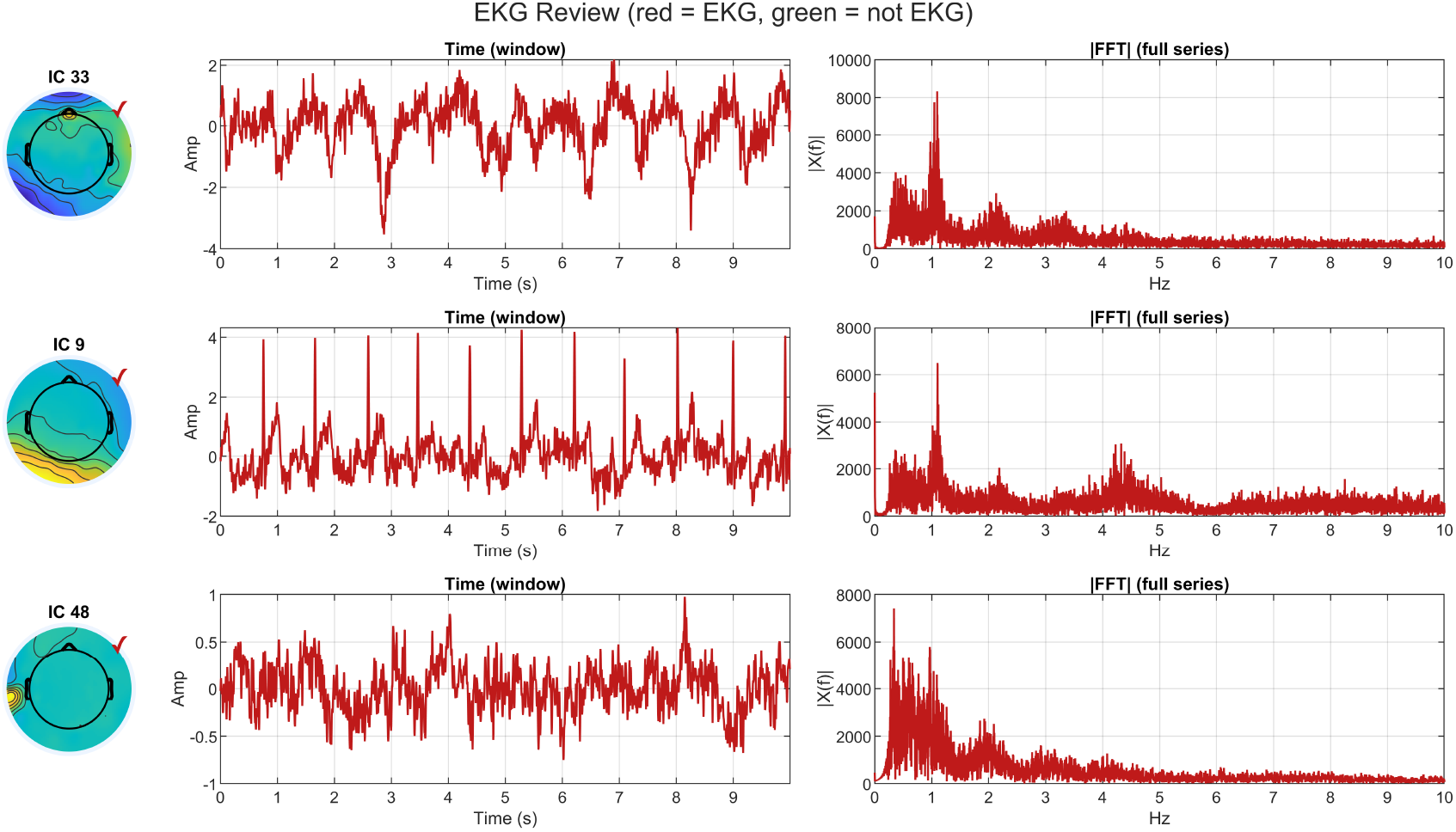
EKG interactive review interface before user confirmation. Components 9, 33, and 48 are initially flagged as EKG candidates by the automated detector.

**Fig. 5.**
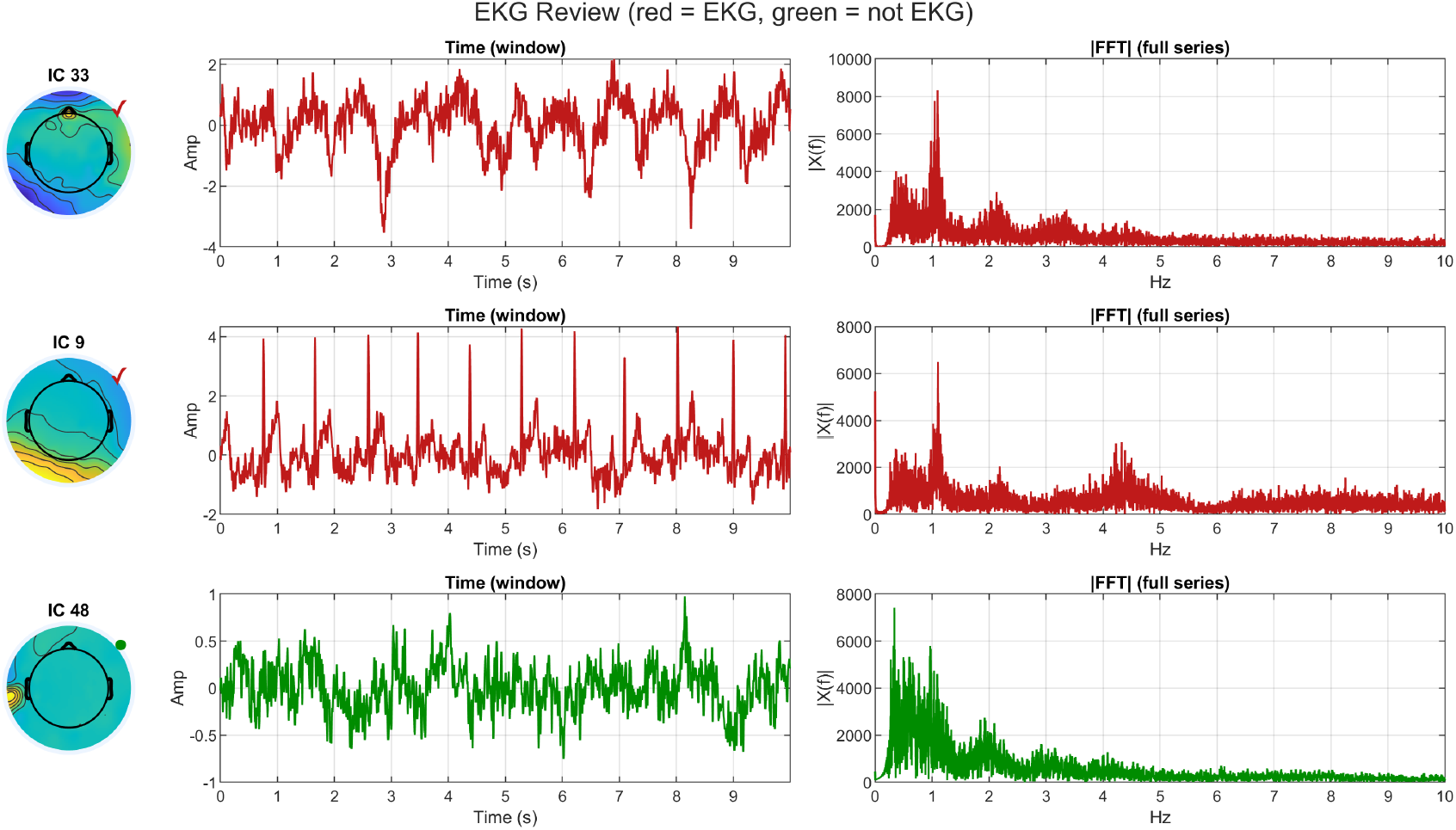
EKG interactive review interface after user confirmation. Components 9 and 33 remain marked for removal, while Component 48 is retained.

In both interfaces, the color coding is intentionally simple: red denotes components currently marked for removal, and green denotes components currently marked to be kept. Clicking on component’s plots toggles its current state, and pressing the *Done* button provided in each pop-up window finalizes the review and returns the updated list of artefactual components (for that artifact family) to MATLAB. If figure saving is enabled, the final UI state is also saved at this stage. This design keeps the review process focused on rapid confirmation of a small number of candidates rather than on exhaustive manual browsing of all ICA components.

### Artifact removal and reconstruction

Once the final component lists have been confirmed, the script combines all rejected sources into a single vector, sets those components’ time courses to zero in ICA space, and reconstructs the cleaned sensor-space signal by inverse projection:

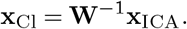

Further elaboration and interpretations for this procedure are provided in the User Manual.

## Discussion

We have presented the first release of a semi-automated Module for identifying cardiac and trigger-related artifact components in ICA-decomposed EEG. The central contribution of this work is a practical and interpretable workflow that makes the review of ICA components faster, more complete, and easier to document. Instead of requiring users to scroll through many components and make ad hoc decisions, the Module ranks a set of candidate components using artifact-specific spectral criteria and then presents those candidates in dedicated review interfaces.

A key design decision in this release was to use frequency-domain structure as the primary signal for automated candidate ranking. For EKG, the harmonic series around a physiologically plausible heart-rate fundamental provided a more stable basis for automated detection than direct peak finding in the time domain, which can be affected by beat-to-beat variability and changes in waveform morphology. For DIN-related contamination, the defining feature was even more strongly spectral. In both cases, the development process suggested that time-domain and topographic cues remain valuable for interpretation, but are less reliable as stand-alone automated rules. This is why the final workflow retains topography and time-domain views in the review interface while relying primarily on spectral metrics to prioritize candidates.

The Module should be viewed as a semi-automated review tool rather than a fully automatic classifier. This distinction matters because the automated score is intended to prioritize candidates, not to replace interpretation. Some components show a clear artifact pattern, whereas others require the user to consider the scalp map, time-domain activity, and spectral structure together before making the final decision. The review interfaces keep the user in control by displaying these three views side by side, so that the final decision reflects the combined evidence. Saving the final review figures also makes the workflow more transparent and easier to document. In this way, the Module improves reproducibility both by reducing the number of components that need inspection and by making the final decisions explicit.

We note that the artifacts addressed in this Module — particularly strictly periodic DIN artifacts — can introduce spectral overlap with periodic EEG responses, such as steady-state evoked potentials (SS-EPs). It is unlikely that a genuine SS-EP will exhibit the highly localized topography and comb-like harmonic structure of the DIN artifact, and ICA will likely separate periodic artifacts and neural activity into separate components. However, users who employ periodic DIN triggering in steady-state paradigms are advised to select DIN and stimulation frequencies with minimal overlapping harmonics in order to avoid inadvertently removing neural responses alongside DIN artifacts.

Several limitations of the current release should be noted. First, the Module depends strongly on the quality of the original data as well as preceding preprocessing steps, particularly filtering, bad-channel handling, and ICA decomposition. Poor ICA separation will necessarily reduce the interpretability of both the automated scores and the review displays. Second, the two detectors differ in generality. EKG contamination is broadly relevant across EEG workflows, whereas DIN-related contamination is system- and setup-dependent and may be absent in many datasets. The DIN detector is therefore most useful in recordings where digital trigger lines are used and DINs are presented at a known repetition period. Third, the current parameter settings and review window lengths were chosen to work well for the provided example recordings, but some adaptation may be needed when sampling rates, frequency resolution, or acquisition characteristics differ substantially from those used here. Finally, the present release is a methods-and-software contribution illustrated on a limited set of example data rather than a comparative benchmark study across multiple datasets, acquisition systems, and laboratories.

These limitations also suggest several directions for future work. A natural next step would be broader validation across recordings with different montages, preprocessing pipelines, and acquisition hardware. On the algorithmic side, future versions could explore adaptive parameter selection. In its present form, the Module provides a practical and transparent workflow for two artifact classes that commonly create manual bottlenecks during post-ICA EEG cleaning.

## ACKNOWLEDGEMENTS

The authors thank members of SENSI and collaborating labs at Stanford University for their feedback and early testing of the Module.

This preprint was prepared using the *HenriquesLab bioRxiv* Overleaf template by Ricardo Henriques.^3^

## AUTHOR CONTRIBUTIONS

Designed the algorithm and implemented the code: AJM. Validated and integrated into larger *SENSI EEG PREPROC* pipelines: AJM, BK. Documented the Module: AJM. Created illustrative analyses: AJM. Wrote the preprint: AJM. Provided supervision, feedback, and editing: BK.

## DECLARATION ON THE USAGE OF AI

The authors used OpenAI’s GPT-5 and Google’s gemini-2.5-flash models for assistance with documentation formatting, function summaries, LaTeX structure, and for brainstorming and exploring conceptual ideas. Additionally, these tools were used to improve flow, word choice, and structure of the User Manual and preprint. No AI tools were used for data analysis, algorithm development, evaluation, or initial drafts of the User Manual or preprint.

## Footnotes

1 https://github.com/edneuro/SENSI-EEG-Preproc-ICA-EKG-Trigger

2 https://purl.stanford.edu/xd818jt7842

3 https://www.overleaf.com/latex/templates/henriqueslab-biorxiv-template/nyprsybwffws

## References

1. T.-P. Jung, S. Makeig, C. Humphries, T.-W. Lee, M. J. McKeown, V. Iragui, and T. J. Sejnowski. Removing electroencephalographic artifacts by blind source separation. Psychophysiology, 2000.

2. Arnaud Delorme and Scott Makeig. EEGLAB: an open source toolbox for analysis of singletrial EEG dynamics including independent component analysis. Journal of Neuroscience Methods, 134(1):9–21, 2004. doi: 10.1016/j.jneumeth.2003.10.009.

3. Maël Chaumon, Dorothy V. M. Bishop, and Nicolas A. Busch. A practical guide to the selection of independent components of the electroencephalogram for artifact correction. Journal of Neuroscience Methods, 250:47–63, 2015. doi: 10.1016/j.jneumeth.2015.02.025.

4. Filipa C. Viola, Jeremy Thorne, Barrie Edmonds, Till Schneider, Tom Eichele, and Stefan Debener. Semi-automatic identification of independent components representing eeg artifact. Clinical Neurophysiology, 120(5):868–877, 2009. doi: 10.1016/j.clinph.2009.01.015.

5. Luca Pion-Tonachini, Kenneth Kreutz-Delgado, and Scott Makeig. Iclabel: An automated electroencephalographic independent component classifier, dataset, and website. NeuroImage, 198:181–197, 2019. doi: 10.1016/j.neuroimage.2019.05.026.

6. Jorge Iriarte, Elena Urrestarazu, Mario Valencia, Manuel Alegre, Alfonso Malanda, Carlos Viteri, and Julio Artieda. Independent component analysis as a tool to eliminate artifacts in eeg: a quantitative study. Journal of Clinical Neurophysiology, 20(4):249–257, 2003. doi: 10.1097/00004691-200307000-00004.

7. Peter J. Allen, Giorgio Polizzi, Keith Krakow, David R. Fish, and Louis Lemieux. Identification of eeg events in the mr scanner: the problem of pulse artifact and a method for its subtraction. NeuroImage, 8(3):229–239, 1998. doi: 10.1006/nimg.1998.0361.

8. Rami K. Niazy, Christian F. Beckmann, Gian Domenico Iannetti, Michael Brady, and Stephen M. Smith. Removal of fmri environment artifacts from eeg data using optimal basis sets. NeuroImage, 28(3):720–737, 2005. doi: 10.1016/j.neuroimage.2005.06.067.

9. Jürgen Dammers, Michael Schiek, Frank Boers, Carmen Silex, Mikhail Zvyagintsev, Uwe Pietrzyk, and Klaus Mathiak. Integration of amplitude and phase statistics for complete artifact removal in independent components of neuromagnetic recordings. IEEE Transactions on Biomedical Engineering, 55(10):2353–2362, 2008. doi: 10.1109/TBME.2008.926677.

10. Magstim EGI Knowledge Center. Synchronizing different types of stimulus events with net station. https://www.egi.com/knowledge-center/item/75-synchronizing-different-types-of-stimulus-events-with-net-station, DINs described as TTL-style 5V signals connected to Net Amps digital inputs.

11. Amilcar J. Malave and Blair Kaneshiro. Sensi-eeg-preproc-ica-ekg-trigger (v1.0): A matlab framework for semi-automated identification of ekg and trigger artifacts in eeg using ica and spectral characteristics, 2026. Version 1.0. DOI: 10.25740/pm515wq0875. Available at: https://github.com/edneuro/ SENSI-EEG-Preproc-ICA-EKG-Trigger.

12. Amilcar J. Malave and Blair Kaneshiro. Example data for the sensi-eeg-preproc-icaekg-trigger module. In Stanford Digital Repository, 2026. Available at: https://purl.stanford.edu/xd818jt7842; accessed 2025-12-17. 10.25740/xd818jt7842.

13. Blair Kaneshiro, Duc T. Nguyen, Anthony M. Norcia, Jacek P. Dmochowski, and Jonathan Berger. Natural music evokes correlated EEG responses reflecting temporal structure and beat. NeuroImage, 214:116559, 2020. ISSN 1053-8119. doi: 10.1016/j.neuroimage.2020.116559.

14. G. Dirlich, L. Vogl, M. Plaschke, and F. Strian. Cardiac field effects on the eeg. Electroencephalography and Clinical Neurophysiology, 102(4):307–315, 1997.

